# Exploring the Relationship between Abundance and Temperature with a Chemostat Model

**DOI:** 10.1101/054940

**Authors:** Cristian A. Solari, Vanina J. Galzenati, Brian J. McGill

## Abstract

Although there is a well developed theory on the relationship between the intrinsic growth rate *r* and temperature *T*, it is not yet clear how *r* relates to abundance, and how abundance relates to *T*. Many species often have stable enough population dynamics that one can talk about a stochastic equilibrium population size *N*\*. There is sometimes an assumption that *N*\* and *r* are positively correlated, but there is lack of evidence for this. To try to understand the relationship between *r*, N*, and *T* we used a simple chemostat model. The model shows that N* not only depends on *r*, but also on the mortality rate, the half-saturation constant of the nutrient limiting *r*, and the conversion coefficient of the limiting nutrient. Our analysis shows that N* positively correlates to r only with high mortality rate and half-saturation constant values. The response curve of N* vs. *T* can be flat, Gaussian, convex, and even temperature independent depending on the values of the variables in the model and their relationship to *T*. Moreover, whenever the populations have not reached equilibrium and might be in the process of doing so, it could be wrongly concluded that N* and *r* are positively correlated. Because of their low half-saturation constants, unless conditions are oligotrophic, microorganisms would tend to have flat abundance response curves to temperature even with high mortality rates. In contrast, unless conditions are eutrophic, it should be easier to get a Gaussian temperature response curve for multicellular organisms because of their high half-saturation constant. This work sheds light to why it is so difficult for any general principles to emerge on the abundance response to temperature. We conclude that directly relating N* to *r* is an oversimplification that should be avoided.

## Introduction

The metabolic rate of any organism basically depends on the concentration of resources in the environment, on the flux of these resources into the organism, on its body size, and on temperature, which determines the rate of biochemical processes (Gillooly, *et al.* 2001). For enzyme-mediated reactions, reaction rates increase from low to high temperature, reaching a maximum, and then rapidly decreasing often due to protein denaturation (Kingsolver, 2009). As in the reaction rates, the response curve of the population intrinsic rate of increase *r* to temperature *T* is generally asymmetric, with a sharper drop off to high temperatures from the optimum (Figure 1). This relationship has been studied in detail and summarized in recent review papers (Huey & Berrigan, 2001; Frazier, *et al.* 2006; Kingsolver & Huey, 2008; Martin & Huey, 2008; Kingsolver, 2009).

Although there is a well developed theory and a robust understanding of the relationship of *r* vs. *T*, it is not yet clear how *r* relates to abundance (or more precisely density in units of individuals per area or volume; McGill, 2006) 2006), and how abundance relates to *T*. There is sometimes an assumption that *r* and abundance are positively correlated, but there is lack of hard evidence for this.

Some species are not even close of having equilibrium population dynamics. In those cases, the idea of abundance may not be well defined and a theory of *r* vs. *T* might be the most useful. Nonetheless, many species do have stable enough population dynamics to meaningfully talk about a stochastic equilibrium population size *N*\*. It is not yet clear how the physiological predictions about *r* translate into predictions about *N*\*.

Given the obvious importance of developing a theory about the effect of temperature on the abundance of species, several routes have been used to attack the question. These include experimental temperature manipulations of communities in natural ecosystems (e.g., Chapin & Shaver, 1985; Suttle, *et al.* 2007), laboratory microcosms (e.g., Davis, *et al.* 1998; Petchey, *et al.* 1999; Jiang & Morin, 2004), observing natural climate change over a few decades in natural communities (e.g., (Kimball, *et al.* 2009), and theoretical population dynamics (e.g., Ives & Gilchrist, 1993; Ives, 1995; Vasseur & McCann, 2005) just to name a few. Nevertheless, from these studies no general principles on the effect of temperature on populations have yet emerged.

**Fig. 1.**
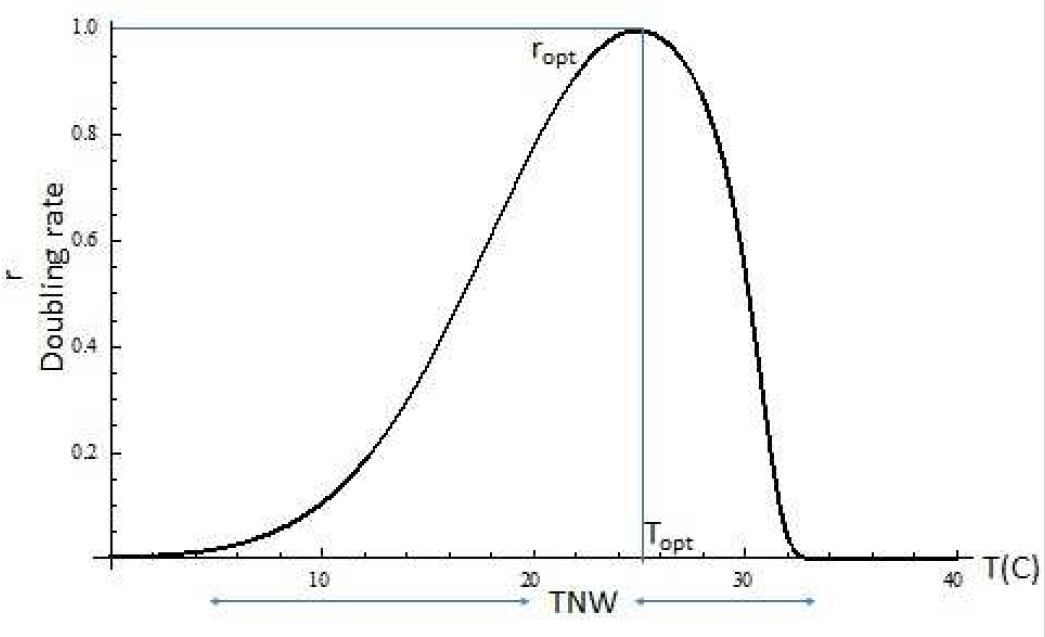
*r* vs. *T*. A typical response curve of the population growth rate *r* to temperature *T* in centigrades (C), where *r_opt_* is the maximal *r* at the optimal temperature *T_opt_*. TNW is the temperature niche width.

For example, Kimball, *et al.* (2009) found that after two decades of natural warming, surprisingly, the cold-adapted annuals increased while the warm-adapted annuals decreased in abundance. In a microcosm experiment with ciliates, Jiang and Morin,(2004) found that temperature had no effect on *N*\* for one species but decreased *N*\* for another in monoculture, the negatively affected species actually competitively excluding the unaffected one at intermediate temperatures. Vasseur and McCann, (2005) predicted that population cycles would be more common and resource biomass would decrease. In another theoretical development (Ives & Gilchrist, 1993; (Ives, 1995) it was predicted that density dependence could buffer/dampen (in intraspecific competition and predator-prey dynamics) or magnify (in interspecific competition) the effect of *T* on *N*\*.

In short, the hypothesis about the performance curve of *r* has received enormous attention, but there has been limited development of theory on the effect of *T* on *N*\*. Gause, (1932, 1934) suggested that *N*\* had a response to *T* similar to that of *r*, a Gaussian bell response curve, but this has received little follow up work. He presented as evidence two field studies done along environmental gradients (in grasshoppers and starfish) and two examples of laboratory experiments where only temperature varied (in the yeast *Sacchromyces* and in *Monia,* a Cladoceran). Later work on flour beetles (*Tribolium;* Birch, 1953; Park, 1954) also showed that the equilibrium population size varied in a modal fashion with temperature, but the number and range of temperatures was not enough to determine the shape in detail.

The simplest approach to explore the relationship between the population growth rate *r*, abundance N* and temperature *T* is to use the chemostat model. A chemostat is a system used in microbiology in which fresh medium is continuously added, while the culture medium is continuously removed at the same rate to keep the volume constant in a dynamical equilibrium. Here, we analyze the chemostat dynamics to try to understand the relationship between *r*, *N*\*, and *T.* We also explore how the abundance response to temperature might differ between a unicellular organism and a more complex multicellular one with germ-soma differentiation.

## The Model

In a chemostat the flow rate *ω* depends on the total volume *V* of the container and the flow *F* in and out of it, *ω* = *F*/*V* In the absence of organisms, the substrate concentration *S* in the container follows, 
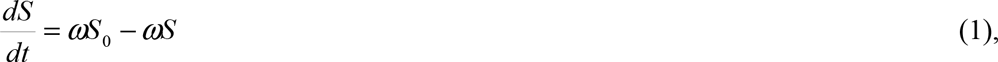
where *S*_*0*_ is the substrate concentration in the medium flowing in and *S*(*t*) is the substrate concentration in the container (Hoppensteadt, 2011). If we use the Monod model to add organisms into the chemostat system (e.g., Droop, 1982), then, 
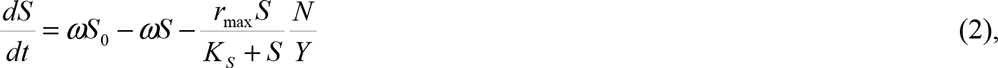

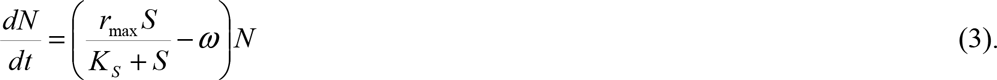
(Equation 2 describes the change in concentration of the limiting substrate *S* due to the inflow of fresh medium (*ωS*_o_; *ω* = flow rate), minus the outflow (*ω*S), minus the substrate consumed by the organisms *N* ([*rS*/(K_S_+*S*] *N/Y*). The substrate consumption depends on the population growth rate *r_max_* when there are no substrate limitations, on the half-saturation constant *K_S_*, which determines how sensitive the organism's growth rate is to substrate limitation, and the substrate conversion coefficient *Y*. (Equation 3 describes the change in population, which depends on *r_max_*, *K_S_*, and the rate *ω* at which the population is discarded (i.e., the mortality rate).

There are two possible equilibrium states in this system (*dN*/*dt* = 0); either when *N* = 0 (the population goes extinct) or when [*rS*/(*K_S_+S*)]−*ω* = 0. If *r* < *ω*, then *N*→0, but if *r* > *ω*,then, 
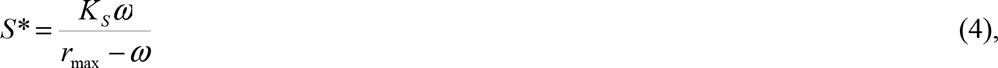

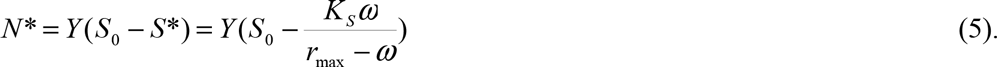
Thus, the equilibrium population *N*\* (i.e., abundance) in a chemostat depends on the total amount of limiting substrate in the system *S*_*o*_, on the flow rate of the system *ω* (i.e., the coupled mortality and nutrient recycling rates), the population growth rate *r*_*max*_, the half saturation constant *K*_*S*_, and the conversion factor *Y* As explained above, it is well known how *r*_*max*_ changes with *T*, but it is not yet clear how *K_s_*, *ω*, and *Y* change as a function of *T*, nor how these variables change as a function of the size and complexity of the organism.

Does abundance *N*\* positively correlates to the population growth rate *r_max_*? In a chemostat it really depends on how much the limiting nutrient negatively affects the population growth rate of the organism in question (i.e., the half-saturation constant *K_s_*), and on its mortality rate *ω* (Figure 2). If *K_s_* is low compared to the amount of limiting substrate in the system (*S*_*o*_; low *K_S_/S_o_* ratio) then the relationship between *N*\* and *r_max_* is flat for most of the *r_max_* values regardless of the mortality rate, with a sharp drop-off of *N*\* at the lower end of *r_max_*. As the *K_S_/S_o_* ratio increases, abundance decreases, but the positive correlation between *N*\* and *r_max_* increases. Increasing the mortality rate *ω* (increasing the *ω*/*r_max_* ratio), also lowers abundance and limits the range of *r_max_* values in which *N*\* is positive, but further accentuates the positive correlation between *N*\* and *r_max_*. In short, it is only with a high negative effect of the limiting nutrient on the population growth rate (i.e., a high *K_S_/S_o_* ratio) and/or a high mortality rate (i.e., a high *ω*/*r_max_* ratio) that we find a significant positive relationship between abundance and the population growth rate.

**Figure 2.**
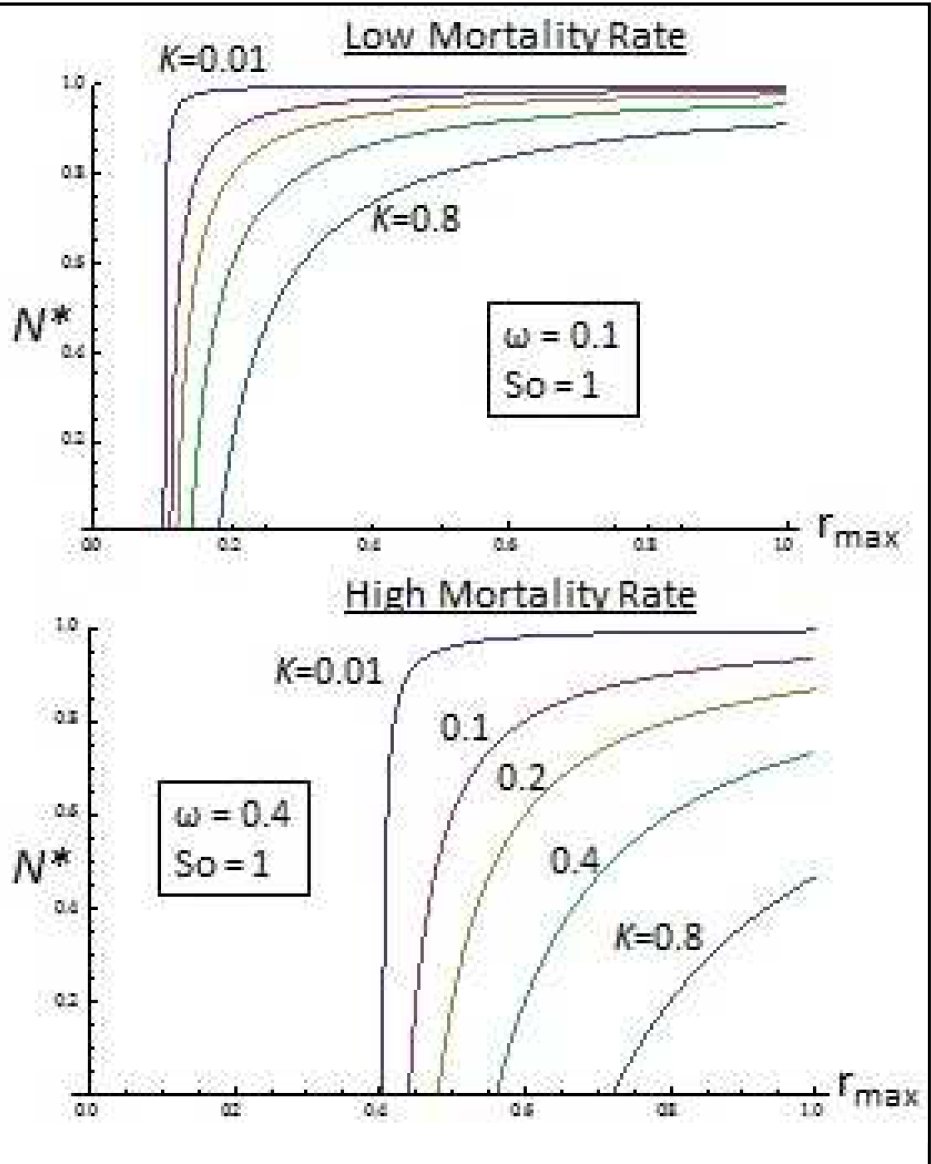
*N*\* vs *r_max_*. The equilibrium population size *N*\* in a chemostat as a function of population growth rate *r_max_* for different values of half-saturation constants *K_S_* and mortality rates *ω*(Equation 5; *Y* and *S_o_* = 1). For low *K_s_* and *ω* values, there is no significant rela ionship between N* and *r_max_*. As both variables increase there is a wider range of values where there is a significant positive correlation between N* and *r_max_*.

Now let's assume *r_max_* has a temperature response curve as described above (Figure 1). To analyze *N*\* vs. *T*, we can use a Gaussian times a Gompertz function to accommodate the nonlinear nature of the relationship between *r_max_* and *T* as described by Frazier, *et al.* (2006),

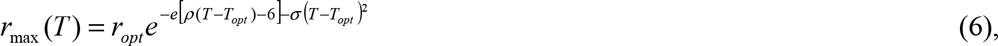
where *r_opt_* is the maximal growth rate at optimal temperature *T_opt_*, ρ represents the increasing part of the population growth rate curve, and σ represents the declining part of the curve. Eq. 5 becomes 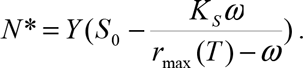

We assume, for now, no relationship between *K_s_*, *ω*, *Y* and *T*. As in *N*\* vs. *r_max_*, we observe that the response curve of *N*\* to *T* is highly dependent on *K_s_* and *ω* levels (Figure 3). As shown in Figure 2, only with high *Ks/S_o_* and *ω*/*r_opt_* ratios we observe a Gaussian temperature response curve for *N*\*. If the mortality rate is low compared to the optimal population growth rate (low ω/*r_opt_* ratio), the equilibrium population (i.e., abundance) response curves to temperature are flat with a steep decline at the edges of the temperature niche width (Figure 3B). Increasing the negative effect of the limiting nutrient on the population growth rate (high *K_s_/S_o_* ratio) slightly decreases the temperature niche width and the population size, but does not change significantly the shape of the response curve. As the mortality rate increases (high *ω*/*r_opt_* ratio), the equilibrium population response curves to temperature become more Gaussian and concave (Figure 3C). In this case, the equilibrium population size and the temperature niche width decrease significantly with the mortality rate.

**Figure 3.**
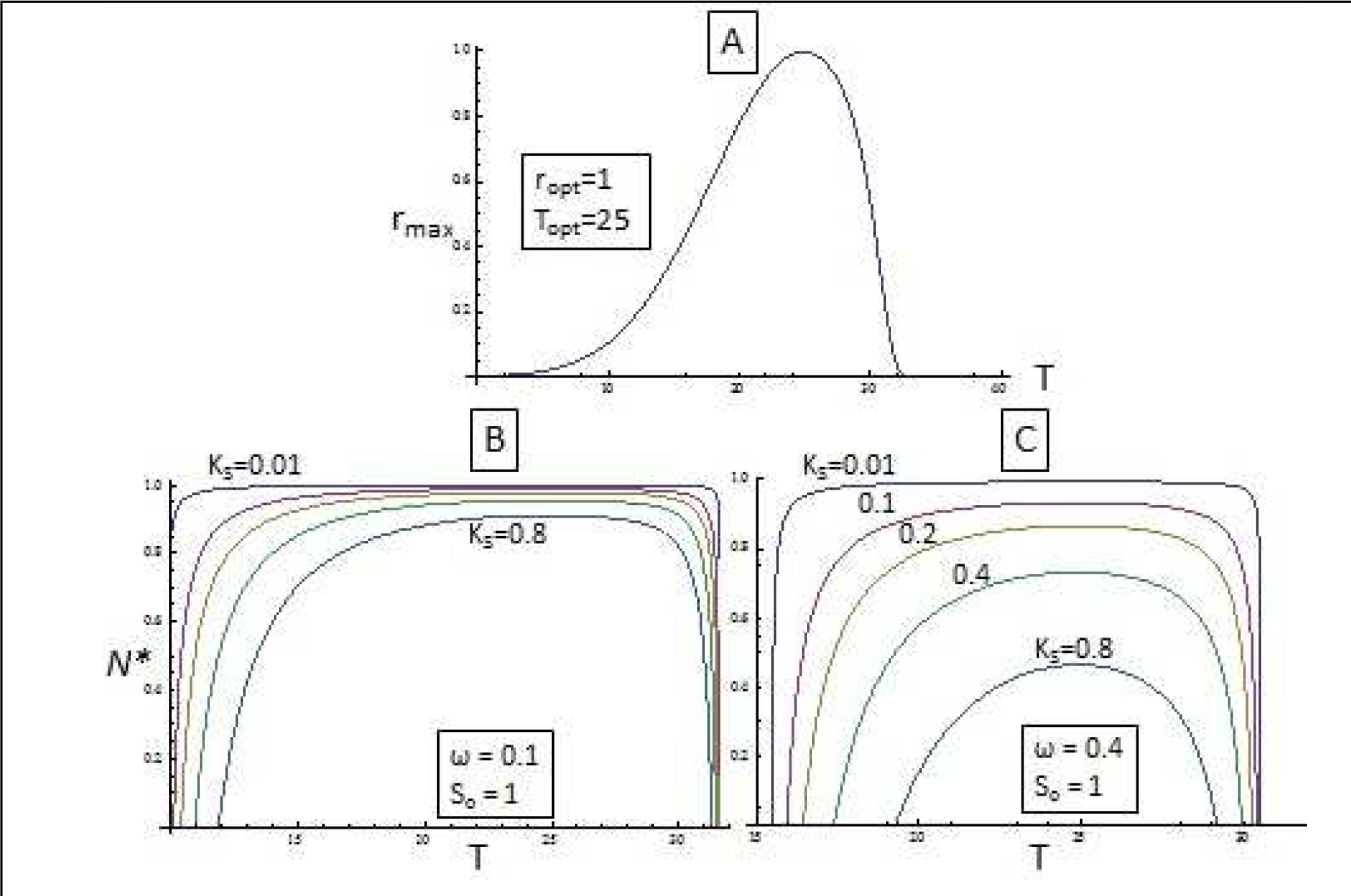
A - *r_max_* vs. *T*(Eq. 6; ρ = 1, ρ = 0.01). B - *N*\* vs. *T*(Eq. 5 with Eq. 6 inserted) with low mortality *ω* for different values of the half-saturation constant *K_s_.*. C - *N*\* vs. *T*(Eq. 5 with Eq. 6 inserted) with high *ω* for different values of *K_s_*. The figure shows that the equilibrium population size has a Gaussian response curve only with high *K_s_* and *ω* values.

What do we know about the relationship between the half-saturation constant *K_S_* and *T*? There has been no consistent pattern observed for the variation of *K_S_* vs. *T* in diverse microorganisms such as algae and bacteria, and for limiting nutrients such as silicate, nitrate, ammonium, and phosphorous (e.g., Mechling and Kilham, 1982; Nedwell, 1999; Tilman, *et al.* 1981). The only consistent pattern found in microorganisms is that *K_S_* values are in general relatively low; meaning that microorganisms can still grow at maximal rates even at very low concentrations of the limiting nutrient. On the other hand, there is evidence that multicellular organisms with germ-soma differentiation have much higher *K_S_* values than unicellular ones, presumably due to the additional nutrients needed to maintain the somatic tissue (e.g., *Volvox sp.;*Senft, *et al.* 1981;). In short, unicellular organisms in general have high population growth rates and low half-saturation constants, but larger multicellular organisms with cellular differentiation have lower population growth rates - due to size/allometric constraints - and higher saturation constants.

In addition, there is evidence that in multicellular *Volvox sp. K_S_* also has a Gaussian temperature response curve similar to that of *r_max_* vs. *T* (Senft, *et al.* 1981, own observations in *Volvox carteri*). This is presumably because the metabolic rate of somatic cells would follow the same temperature response curve of reproductive cells, increasing the metabolic need of soma at optimal temperatures and decreasing it at suboptimal ones. To analyze this, for the sake of simplicity, we assumed for the hypothetical multicellular organism that *K_S_* has the same response curve as *r_max_(T)* (Eq. 6), *K_S_*(*T*) = *K*_*max*_r_*max*_(*T*), where *K_max_* is the maximum half-saturation constant value at the optimal temperature (for *K_S_(T)* we use the same parameters as in *r_max_(T)* in Eq. 6). Thus, Eq. 5 becomes 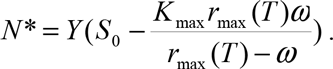

Figure 4 shows *N*\* vs. *T* for a hypothetical unicellular organism with a low *K_S_/S_o_* ratio and a constant *K_S_*, and for a multicellular one with *K_S_*(*T*) = *K*_*max*_*r*_*max*_(*T*) and a high *K_max_/S_o_* ratio.

From a nutrient availability point of view, if conditions are oligotrophic (low *S_o_*), then a unicellular population could also have a high *K_S_*/*S_o_* ratio, thus, the *N*\* response to *T* could be similar to that of a population of multicellular organisms. On the other hand, if the conditions are eutrophic (high *S_o_*), then the *K_max_/S_o_* ratio for the multicellular population would also be low and its response similar to that of the unicellular population. In short, unicellular or multicellular populations in eutrophic conditions would have a similar response curve to *T*, and unicellular populations in oligotrophic conditions would have a response curve similar to that of multicellular populations.

**Figure 4.**
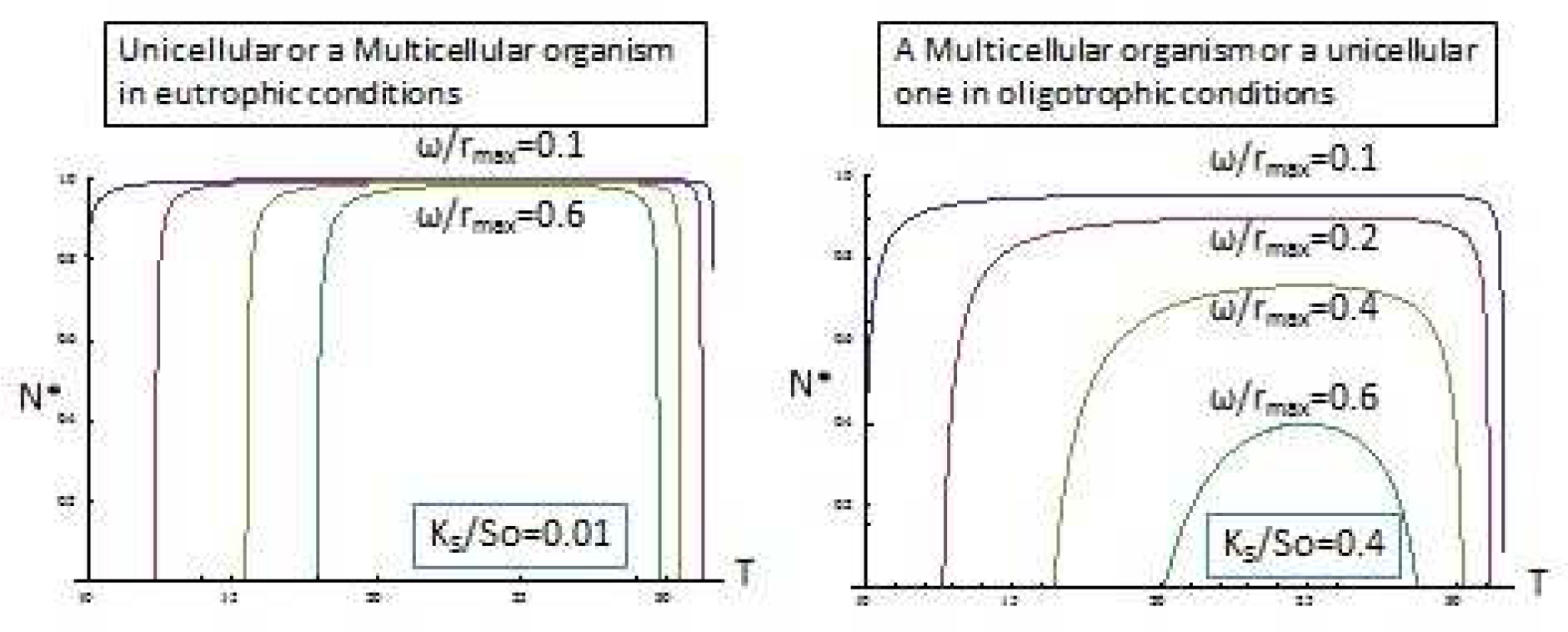
*N*\* vs. *T* for hypothetical unicellular and multicellular organisms in eutrophic and oligotrophic conditions (*r_opt_*, *Y*, and *S_o_* = 1; ρ = 1, σ = 0.01). For the eutrophic condition case, curves are always flat; increasing the mortality rate slightly decreases *N*\*, but greatly decreases the temperature niche width. For the oligotrophic-unicellular/multicellular case, curves are flat for low mortality rates, but as the mortality rate increases the response curve becomes more concave and both *N*\* and the temperature niche width decrease significantly.

In the hypothetical unicellular/eutrophic case the equilibrium population (i.e., abundance) response curves to temperature are flat with a steep decline at the suboptimal high and low temperatures (Figure 5). Increasing the mortality rate decreases the temperature niche width, but does not change significantly the shape of the response curve nor the population size in optimal temperatures. On the other hand, in the multicellular/oligotrophic case the equilibrium population response curves to temperature become more Gaussian and concave as the mortality rate increases. In this case, both the equilibrium population size and the temperature niche width decrease significantly with the mortality rate.

What about the relationship between the mortality rate *ω* and *T*? In a chemostat *ω* is temperature independent because it is artificially adjusted by changing the flow/removal rate in the system. But what if we envision a community of ectotherms such as a plankton community, where we are tracking the abundance of the phytoplankton? The predation rate can be the mainfactor affecting the mortality rate, and the zooplankton grazers might have metabolic and feeding rates that respond to temperature in the same way the phytoplankton population growth rates do. For example, it was shown that copepods that graze on phytoplankton have a feeding rate that follows a domeshaped patte*ω* as a function of temperature (e.g., Almeda, *et al.* 2013; Garrido, *et al.* 2013; Moller, *et al.* 2012).

**Figure 5.**
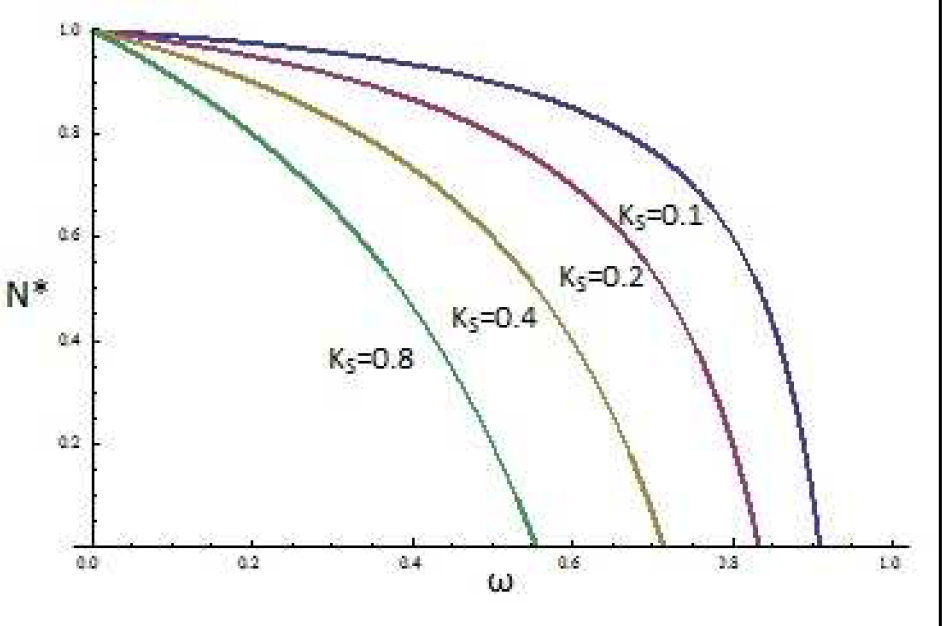
*N*\* vs. when*ω*(*T*) = *ω*_max_*r*_max_ (*T*) for different values of *K_S_*. (*S_o_* and *Y* = 1). *N*\* decreases as *ω* increases; its decrease is more pronounced with higher *K_S_* values.

If again for the sake of simplicity we assume that *ω* has the same response curve as *r_max_*(*T*) (Eq. 1), then *ω*(*T*) = *ω*_max_*r*_max_(*T*), where *ω*_max_ is the maximum mortality rate (i.e., predation) at the optimal temperature. If we use the same parameter values for *r*(*T*) and *ω*(*T*), then the temperature term cancels out and Eq. 5 becomes temperature independent;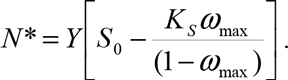

In this case, *N*\* only depends on *K_s_* and *ω_max_*. Figure 5 shows how *N*\* decreases as *ω* increases; its decrease is more pronounced with higher *K_s_* values.

If we now assume that both *K_S_*(*T*) and *ω*(*T*) have the same response curve as *r_max_*(*T*), then Eq. 5 becomes 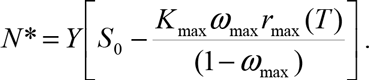

Surprisingly, the temperature response curve flips, becoming convex (Figure 6). The populations of the hypothetical organisms are better off at suboptimal temperatures because both the half-saturation constant and the mortality rate are at their maximum values at optimal temperatures, negatively affecting abundance in the best conditions for population growth.

**Figure 6.**
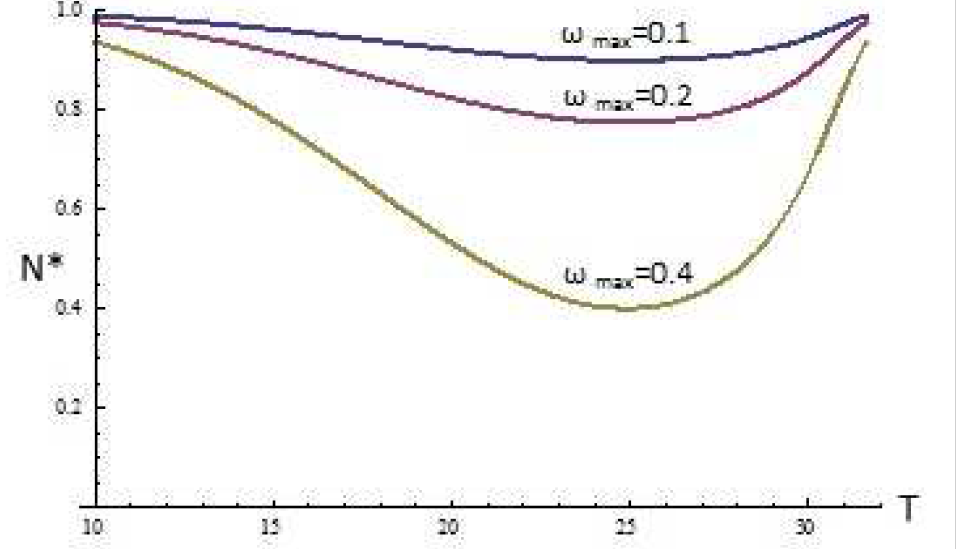
*N*\* vs. *T* for different values of *ω_max_* (ρ = 1, σ = 0.01; *r_opt_*, *Y*, and *S_o_* = 1, *K_max_* = 0.1) for the case where both *K_S_*(*T*) and σ(*T*) have the same response curve as *r_max_*(*T*). The populations of the hypothetical organisms are larger at suboptimal temperatures because both the half-saturation constant and the mortality rate increase at optimal temperatures.

What about the relationship between the conversion coefficient *Y* and *T*? The temperature-size rule proposes that ectotherms that develop at higher temperatures are relatively smaller as adults than when they develop at lower temperatures. There is plenty of evidence for this rule in multiple taxa (Kingsolver and Huey, 2008). Therefore, if we assume abundance to be density in units of individuals per area or volume, the conversion coefficient *Y* may change as temperature increases, since smaller individuals would need fewer nutrients to develop. Some of the relationships reported between size and temperature have an approximately negative linear slope, so for simplicity, we can just assume a positive linear relationship between *Y* and *T* (*Y*(*T*) = *aT* + b) since less need for nutrients means more individuals per substrate absorbed from the medium. Figure 7 shows how, *N*\* response curves get skewed to higher abundance peaks at higher temperatures as *Y* increases with *T*. What would be an almost flat response curve if the relationship between *Y* and *T* is not taken into account (for *ω* = 0.1, Figure 3) becomes a concave curve with a peak at a higher temperature if the size-temperature rule is taken into account (for *ω* = 0.1, Figure 7).

**Figure 7.**
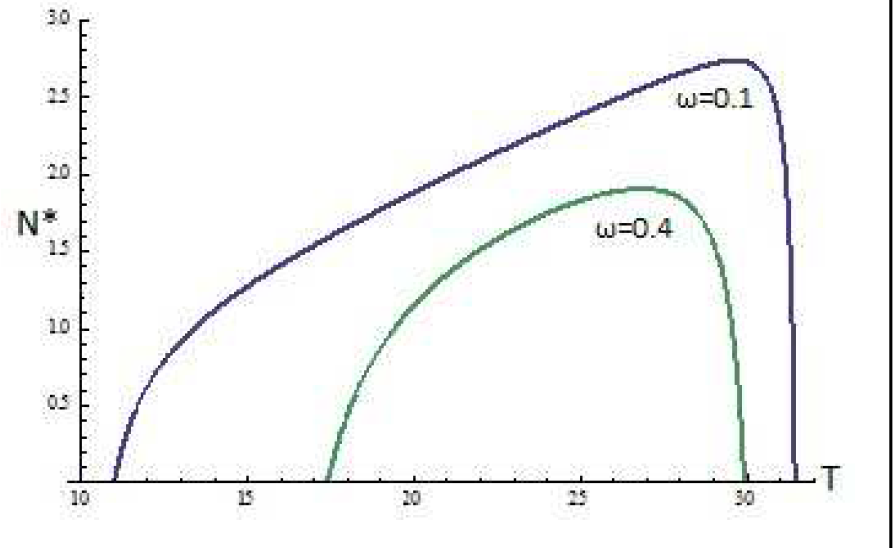
Hotter is smaller. *N*\* vs. *T* for different values of *ω* (ρ = 1, σ = 0.01; *r_opt_* and *S_o_* = 1, K_S_ = 0.4; *Y* = 0.1*T*) for the case where *r_max_*(*T*). *N*\* response curves get skewed to higher abundance peaks at higher temperatures because smaller organisms need less nutrients.

So far we have only analyzed populations that have reached equilibrium population size at different temperatures (i.e., *N*\*, Eq. 5). Although we have showed, for example, that *N*\* does not change significantly with *T* at low *K_S_/S_o_* and *ω*/*r_max_* ratios, populations growing at optimal temperatures with high *r_max_* values would reach equilibrium population size much faster than those growing at suboptimal temperatures. If we solve Eq. 3 for time *t* then, 
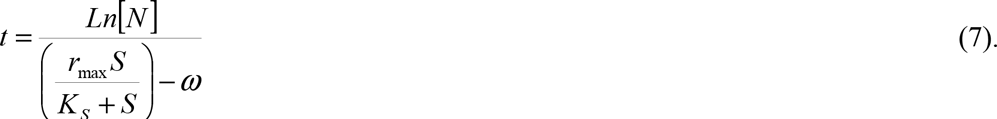
To illustrate this point and for the sake of simplicity we will just assume that *K_s_* ~ 0, thus, *t* ≈ *Ln*[*N*]/(*r*_max_ - *ω*). Figure 8A shows how the time to reach a certain population size increasesexponentially as *r_max_* decreases (Figure 8A). If we plot populations at different time steps before reaching an arbitrary population size 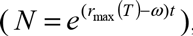, the abundance at those time steps would seem to have a Gaussian bell response curve as a function of *T* instead of the flat response we see when the population reaches equilibrium (Figure 8B). In short, if the population does not reach equilibrium, its abundance temperature response curve could resemble the population growth rate curve.

**Figure 8.**
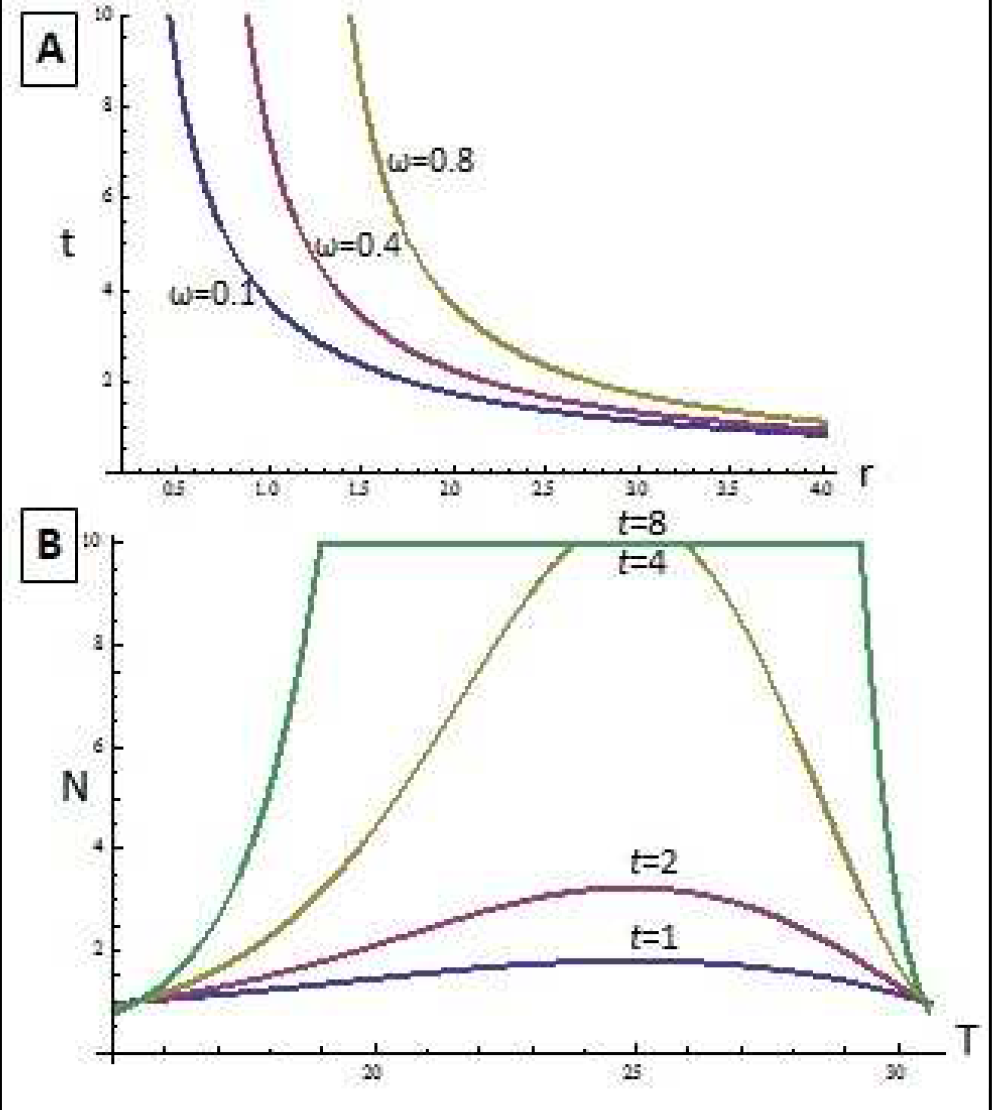
A-*t* vs. *r* for different values of *ω* ((Eq. 7, *K_S_* = 0, *N* = 10). Time to reach a certain population size increases exponentially *as r_max_* decreases. B – *N* vs. *T* for different values of *t* (*K_S_* = 0 and *r*(*T*), (Eq. 6). We arbitrarily assume that maximum population *N* = 10. The abundance at different time steps could seem to have a Gaussian bell response curve as a function of *T* instead of the flat response we see when the population reaches equilibrium

## Discussion

In this paper we have explored the relationship between abundance and temperature using a simple chemostat model. We have shown that the equilibrium population size not only depends on the population growth rate, but also on its mortality rate - and coupled nutrient recycling rate -, and on the relationship the organisms in question have with the limiting nutrient in the system (Eq. 5). Abundance positively correlates to the population growth rate in a chemostat only with a high negative effect of the limiting nutrient on the population growth rate (i.e., a high *K_s_/S_o_* ratio) and/or a high mortality rate compared to the population growth rate (i.e., a high *ω*/*r_max_* ratio; Figure 2).

If we assume a typical population growth rate response curve to temperature and leave all the other variables constant (Figure 1, Eq 6), again only with a high negative effect of the limiting nutrient on the population growth rate and/or a high mortality rate, the equilibrium population response curves to temperature become Gaussian as a function of temperature (Figure 3). With low mortality or half-saturation constant values, the relationship between abundance and temperature remains flat for almost all of the temperature niche width.

In general, unicellular organisms have high population growth rates and low half-saturation constants, but larger multicellular organisms with cellular differentiation have lower population growth rates and higher saturation constants. Moreover, evidence shows that in multicellular organisms the half-saturation constant for a limiting nutrient might have a Gaussian response curve to temperature similar to the population growth rate one (e.g., *Volvox* sp.; Senft, *et al.* 1981; own observations in *Volvox carteri*).

Our analysis shows that a unicellular or a multicellular population in eutrophic conditions could have a similar response curve to temperature (low *K_S_/S_o_* ratio) because the higher half-saturation constant in a multicellular organism due to the metabolic need of the soma would be minimized by the high concentration of nutrients in the system (Figure 4). In this case, even with high mortality rates there is no relationship between abundance and temperature in almost all of the temperature niche width. In contrast, a unicellular population in oligotrophic conditions would have a response curve similar to a multicellular population (high *K_S_/S_o_* ratio) because the low concentration of nutrients in the system would augment the limiting nutrient negative effect on the unicellular population. In this case, there is an increased Gaussian response curve of abundance to temperature with increasing mortality rates.

If the mortality rate has a similar temperature response curve as the population growth rate (e.g., the predation rate temperature response curve is similar to the prey population growth rate response curve), abundance might even become temperature independent, and only depend on the maximum mortality rate and half-saturation constants (Figure 5). Furthermore, if both the mortality rate and the half-saturation constant have the same temperature response curve as the population growth rate (i.e., as in a multicellular population), then the curves flip and become convex, organisms having higher abundance at suboptimal temperatures since the highest mortality and half-saturation constant levels coincide with the optimal temperature for growth (Figure 6). In addition to all these scenarios, the ubiquitous temperature-size rule would increase abundance at higher temperatures since organisms would need fewer nutrients per individual as they develop to a smaller adult size (Figure 7), skewing abundance temperature response curves to higher temperatures.

To summarize, with a simple chemostat model we have shown why abundance might respond to temperature in many ways. The response curve of abundance to temperature can be flat, concave, convex and even temperature independent depending on the population growth and mortality rates, the half-saturation constant, the amount of limiting nutrient in the system, and the conversion coefficient of the nutrient. Even if the system is closed and there is no nutrient concentration change in the flowing medium, the availability of the limiting nutrient to the organisms might change since diffusion coefficients are also temperature dependent, therefore possibly affecting abundance. In conclusion, directly relating abundance to the population growth rate is an oversimplification that should be avoided.

Finally, it is important to point out that if the population of interest has not reached equilibrium and might be in the process of doing so, an observer can reach the wrong conclusion that abundance and population growth rates are positively correlated and have similar response curves for temperature (Figure 8). Because of the limited timeframe of studies, researchers can reach wrong conclusions on how abundance is affected by temperature. If the population is allowed to reach equilibrium, depending on the conditions of the system where the population is growing, there might be no relationship between abundance and growth rate, and between abundance and temperature.

Of course, in natural systems several trophic levels and species interact with one another; temperature, light intensity, and nutrient availability do not remain constant, and the recycling of nutrients and the mortality rate are not directly coupled as they are in a chemostat. We have not taken into account very important population dynamics aspects such as multispecies interactions (e.g., competition for resources; Tilman, *et al.* 1981), organisms adaptation to temperature change (e.g., Thomas, *et al.* 2012; in *Chlorella vulgaris,* Padfield, *et al.* 2015), changes in the nutrient recycling rate due to, for example, the temperature dependence of the detritivores metabolic rate, changes in the total amount of nutrients in the system due to net inflows/outflows from other sources or sinks, just to name a few.

In addition, complexities such as behavioral thermoregulation and water vs. heat balance are not a factor in our model. Terrestrial organisms can behaviorally thermoregulate their body temperatures to deviate significantly from the air. Moreover, water and heat balance are confounded for terrestrial organisms - both plants and animals cool themselves by evaporation, resulting in a strong water-temperature interaction.

Nonetheless, if a population in a chemostat can be used as an oversimplified analogy for a population that is at equilibrium in a stable ecosystem, the model analysis shows why it is so difficult for general principles to emerge on the effect of temperature on populations. When studying populations, it is difficult to know what nutrient is limiting population growth, what the main mortality factor is, what the conversion coefficient is, etc., and how all of these factors are changing with temperature. Are these populations observed at some kind of stochastic equilibrium? Are they on their way to reaching a new one? Or regular perturbation will never allow them to reach one? How fast do these populations adapt to a new temperature regime or a new limiting nutrient? From our very limited analysis, we conclude that it is difficult to make well founded predictions about the outcome of abundance due to temperature change unless all of these factors are well known and taken into account.

In the future we plan to set up population experiments at different temperatures, nutrient concentrations, and mortality rates, in order to measure all the variables in the chemostat model (*r*, *ω*, *N*\*, *K*, *Y*) and check whether the predicted abundance outcomes of the model hold. We believe that laboratory-based microbial model systems to address ecological questions have been, and are still underused (Jessup, *et al.* 2004). Although they are not intended to reproduce natural conditions, their simplicity allows addressing questions that would be inaccessible through other means.

We plan to use the volvocine green algae as a model system (Figure 9). The Volvocales are facultatively sexual, uni-and multicellular, flagellated, photosynthetic, haploid organisms with varying degrees of complexity stemming from differences in colony size, colony structure, and germ-soma specialization. They range from the unicellular *Chlamydomonas* (Fig. 9A), to multicellular individuals comprising 1,000–50,000 cells with complete germ-soma separation, e.g. *Volvox* (Fig. 9E,F; Kirk, 1998; Herron and Michod, 2008; Nozaki, *et al.* 2006; Solari, *et al.* 2006). Due to their range of sizes, they enable the study of scaling laws: the number of cells ranges from 10^0^ *(Chlamydomonas)* to ~10^4^ *(Volvox barberi*). All kinds of communities can be assembled with organisms of different sizes and complexity, but with similar cell biology.

The first set of experiments will consist of populations grown in axenic (sterile) conditions, thus having only one trophic level (producers) in monoculture and competing with each other (polyculture). In the second part of the project we will use non-axenic cultures adding the detritivorous trophic level. We expect the population dynamics of these experiments to be totally different from those of the experiments made with axenic cultures (we continuously observe this in our axenic and nonaxenic stock cultures). We also expect totally different dynamics between unicellular, differentiated, and germ-soma differentiated *Volvox* species, since the large multicellular species will shed more organic material that the bacteria can consume (ECM with somatic cells) than the non-differentiated or unicellular ones.

**Figure 9.**
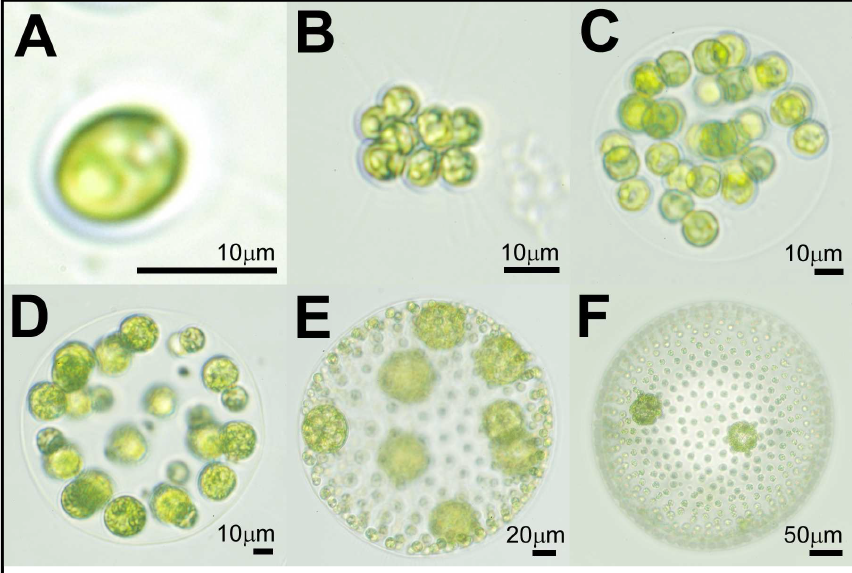
Pictures of various species of Volvocales showing the increase in size and complexity. A-*Chlamydomonas reinhardtii*. B-*Gonium pectorale*. C-*Eudorina elegans*. D-*Pleodorina californica*. E-*Volvox carteri*. F-*Volvox aureus*.

We will then introduce a third trophic level (e.g., the phagotroph Euglenoid *Peranema trichophorum,* which we already keep and use, Solari, *et al.* 2015. The filter-feeding rotifer *Brachionus calyciflorus* and unicellular protist *Paramecium tetraurelia* are possible alternatives for predators). This will be of interest for two reasons. First, it will explore a second type of species interaction (predator/prey or more precisely herbivory). Second, these are size-dependent predators that will greatly tip the competitive balance among the species.

## Acknowledgements

This work was supported in part by CONICET grant PIP 283, Ministry of Science grant PICT 2011–1435, and the Universidad de Buenos Aires.

